# Zinc finger transcription factors *BnaSTOP2s* regulate sulfur metabolism and confer resistance to *Sclerotinia sclerotiorum* in *Brassica napus*

**DOI:** 10.1101/2024.05.17.594441

**Authors:** Lihong Dai, Zhaoqi Xie, Tianxu Ai, Yushun Jiao, Xiaoyi Lian, Angchen Long, Jinyun Zhang, Guangsheng Yang, Dengfeng Hong

## Abstract

Rapeseed (*Brassica napus* L.) has a high sulfur requirement for optimal growth, development, and pathogen resistance. In this study, we identified zinc finger transcription factors, *BnaSTOP2s*, that play key roles in sulfur metabolism and *Sclerotinia sclerotiorum* resistance. First, our results suggested that *BnaSTOP2s* are involved in sulfur as evidenced from extensive protein interaction screening. Knockout of *BnaSTOP2s* reduced the response sensitivity in both sulfur-deficient and sulfur-excessive conditions by promoting the elongation of primary roots of seedlings. Furthermore, the content of essential sulfur-containing metabolites, including glucosinolate and glutathione, were substantially down-regulated in roots and leaves of *Bnastop2* mutants, which is consistent with the significantly lowered transcriptional levels of key players of GSL synthesis and transportation, *BnaMYB28s* and *BnaGTR2s*, respectively. Through comprehensive RNA-seq analysis, we revealed the substantial effect of *BnaSTOP2s* on sulfur metabolism from source to sink. Additionally, we observed a significant decrease while increase in leaf lesion sizes of the *BnaSTOP2*-OE and *Bnastop2* mutants, respectively, when compared to the wild type during *Sclerotinia sclerotiorum* infection, suggesting the vital role of *BnaSTOP2* in plant defense response. Overall, our findings highlight that *BnaSTOP2s* seems to be global regulators of sulfur metabolism and confer resistance to *Sclerotinia sclerotiorum* infection in *B. napus*.

## INTRODUCTION

Sulfur, as a vital nutrient in plants, plays a pivotal role in the biosynthesis of amino acids and proteins, which are indispensable for the intricate processes of plant growth and development (Narayan et al., 2022). Plants can optimize the utilization of available sulfur to meet the demands for higher yield, superior quality, and enhanced stress resistance (Kopriva et al., 2019). The principal source of sulfur in plants is sulfate, which is assimilated from the soil through the roots (sulfur source) and transported by sulfate transporters (SULTRs) to various organs (sulfur sink), such as leaves, siliques and seeds (Hawkesford and De Kok, 2006; Zhao et al., 2008). Upon entering the plastid, sulfate is catalyzed by ATPS (ATP sulfatase) to form APS (adenosine 5-phosphate sulfonic anhydride) due to its instability (Takahashi et al., 2011). APS is further reduced to SO_3_^2-^ by APR, and then reacts with ferriredoxin under the action of sulfite reductase to form S^2-^. Subsequently, under the influence of serine acetyltransferase (SAT) and O-acetylserine (thiol) lyase (OASTL), S^2-^ and O-acetylserine react to produce cysteine (Wirtz et al., 2004), representing a crucial step in the conversion of inorganic sulfur to organic compounds. Additionally, sulfur is present in sulfur-containing cofactors like biotin, thiamine, and coenzyme A, which are vital for enzymatic reactions and metabolic processes in plants.

Moreover, sulfur also actively participates in the synthesis of various secondary metabolites, such as glucosinolate (GSL) in plants belonging to the Brassicaceae family and S-alk(en)yl cysteine sulfoxides in species from the Allium genus, contributing to defense mechanisms and plant adaptation (Francioso et al., 2020). Amongst these compounds, cysteine assumes a pivotal role in the biosynthesis of organic sulfur compounds. It becomes integrated into proteins and the tripeptide glutathione, while concurrently acting as the sulfur-based precursor for the synthesis of the indispensable amino acid methionine (Takahashi et al., 2011). Then there is a wide range of known sulfate metabolites that play various roles in plant resistance to biotic and abiotic stresses (Capaldi et al., 2015), with GSL considered as significant storage forms of sulfur (Aarabi et al., 2020). GSLs and their hydrolysis products such as isothiocyanates (ITCs), nitriles, and thiocyanates, participate in conferring plant resistance against fungal pathogens, particularly *Sclerotinia sclerotiorum* (*S. sclerotiorum*) (Chittem et al., 2020; Hunziker et al., 2020; Sotelo et al., 2015; Stotz et al., 2011), as well as plant innate immunity (Chen et al., 2020; Yang et al., 2020). In summary, sulfur is a vital element for diverse biological processes, playing a pivotal role in plant growth, development, and adaptation to the environment. Its presence in a wide array of compounds, contributes to fortifying defense mechanisms against pathogens.

Rapeseed (*Brassica napus*, AACC, 2n=38) has a high nutrient requirement of sulfur (Grant et al., 2012; Verma et al., 2022). Meanwhile, like Arabidopsis, it also produces abundant GSLs, a high content sulfur-containing secondary metabolites, in roots, leaves and silique walls, a large part of which are destinated to seeds (Petersen et al., 2002; Sanden et al., 2024), though GSLs have been dramatically reduced in canola-type (also call ‘double-low’) rapeseed. As rapeseed has gained importance as one of the main edible plant oil sources worldwide and also an agricultural industrial crop due to its diverse applications (Chao et al., 2017; Wang et al., 2018), an increasing attention should be paid to understand how sulfur mediates the developmental process as well as the yield and quality formation in *B. napus*.

*STOP2* (*Sensitive to Proton Root Toxicity 2*), a Cys(2)His(2)-type zinc finger transcription factor, has garnered relatively less attention compared to its unique homologue *STOP1* (*Sensitive to Proton Root Toxicity 1*), which can directly engage with the promoter of the *NITRATE TRANSPORTER 1.1*. This interaction leads to the activation of transcription in response to low pH values, consequently resulting in the upregulation of nitrate absorption (Ye et al., 2021). Nevertheless, *STOP2*, albeit being a physiologically minor isoform of *STOP1*, possesses the capacity to activate gene expression in a similar manner like *STOP1* (Kobayashi et al., 2014). Here we conducted a comprehensive phenotyping and multi-omics analysis based on the knockout and over-expression mutants of *BnaSTOP2* genes in *B. napus*. According to our data, we have established that *BnaSTOP2*s are global regulators of sulfur metabolism and play a significant role in resistance to *Sclerotinia sclerotiorum* in *B. napus*.

## RESULTS

### Characterization of *BnaSTOP2* homologues in *B. napus*

A comparative analysis showed that STOP2 shares only 34.7% sequence similarity with STOP1 in amino acid sequence in Arabidopsis (Supplemental Figure 1), suggesting the possibility of functional divergence between two members. To explore the evolutionary conservation and functional relevance of *BnaSTOP2,* we conducted a sequence comparison between all the homologous of *STOP2* within *B. napus*. Using STOP2 as a query, we identified six homologous genes according to the sequence similarity in the genome of *B. napus*, and they can be divided into two groups, with BnaA02.1.STOP2 showing a much higher sequence similarity with STOP2 than the other five copies (Figure 1A). All the BnaSTOP2 homologues in *B. napus* encompass two highly conserved zinc finger domains, strategically positioned within the central and C-terminal regions of its amino acid sequence (Supplemental Figure 2A), suggesting a potential functional redundancy among these genes.

**Figure 1.**
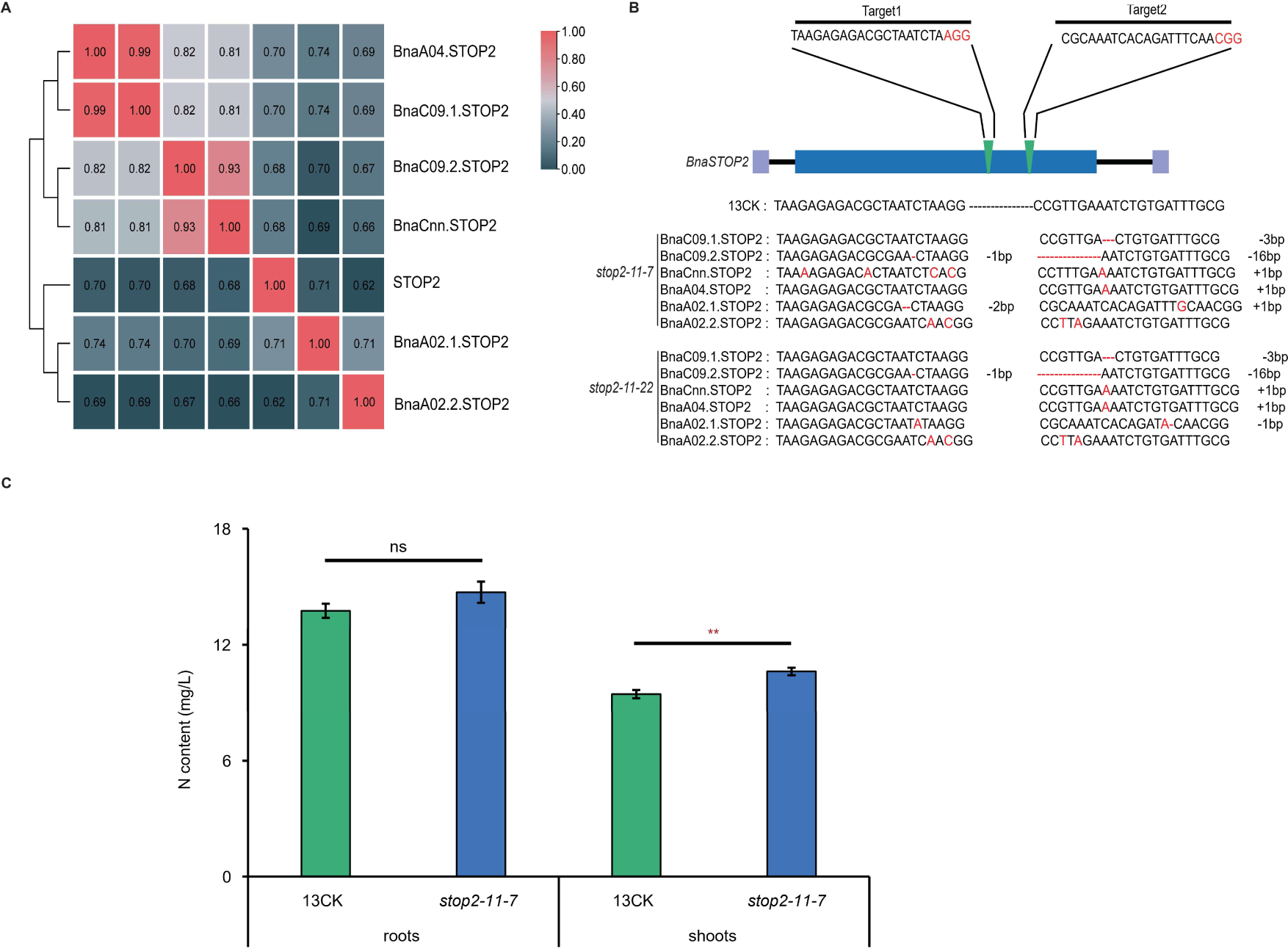
Generation of CRISPR/Cas9-induced mutations in *BnaSTOP2*. (**A**) Amino acid sequence alignment of STOP2 in *Brassica napus* and Arabidopsis. (**B**) The indel situation of the *Bnastop2* knockout site compared to the WT sequence. The sgRNA target sites are marked with arrows, the PAM sequence is colored in red, and the indel mutations are colored in red. (**C**) N content (mg/L) in roots and shoots of *Bnastop2* mutants. Differences between mutants and WT are significant at p< 0.05 (*) and p<0.01 (**) by two-tailed Student’s t test.

To explore the biological function of *BnaSTOP2s*, we used the CRISPR-Cas9 tool to knockout *BnaSTOP2* genes (Supplemental Dataset 1). We obtained a heterozygous sextuple mutant *stop2-11* in the T_0_ generation, from which the offsprings lines, *stop2-11-7* and *stop2-11-22*, emerged as homozygous sextuple mutants in the T_1_ generation (Figure 1B). These mutants exhibited base substitutions, deletions, and insertions that resulted in amino acid alterations, frameshifts, or premature stop codons in different paralogues. Consequently, there were substitutions and truncations in the zinc finger domain at the amino acid level (Supplemental Figure 2). To simplify our investigation and gather more focused insights, we prioritized the sextuple mutant line *stop2-11-7* (designated as *Bnastop2*) for further analysis. Initially, RT-qPCR analyses revealed that *BnaSTOP2* exhibits primary expression in roots, and the expression were markedly down-regulated in both roots and leaves of *Bnastop2* mutants (Supplemental Figure 3, Supplemental Dataset 2). These findings shed light on the tissue-specific regulatory mechanisms associated with *BnaSTOP2* and emphasized its potential role in roots.

Considering that *STOP1* contributes to nitrate uptake and nitrogen metabolism in Arabidopsis (Tokizawa et al., 2023; Ye et al., 2021), we endeavored to unravel the potential involvement of *BnaSTOP2* in nitrogen metabolism. As shown in Figure 1C and Supplemental Dataset 3, though our analyses revealed a reduction of nitrogen content in the shoots of *Bnastop2* mutants, the magnitude of change seemed slight; moreover, no significant variation was observed in the roots. These findings suggested that *BnaSTOP2s* may play other roles more than in influencing nitrogen metabolism in shoots.

### *BnaSTOP2* is involved in regulating sulfur metabolism in *B. napus*

It is noteworthy that Zinc finger structural proteins generally act as transcriptional factors to regulate the gene expression of functional genes (Noman et al., 2019). To discover the potential functions of *BnaSTOP2* genes that contains two conserved Zinc finger domains, we attempted to identify the interaction proteins of it by exploring a yeast two-hybrid (Y2H) library screening. BnaA02.1.STOP2, that shares a protein sequence similarity of 71% with STOP2 (Figure 1A), was used as a bait protein. A comprehensive set of 399 interacting proteins was identified (Supplemental Dataset 4). Further examination of the KEGG database unveiled that these interacting proteins are implicated in metabolism pathways, crucial pathways encompassing oxidative phosphorylation, mRNA surveillance pathway, amino acid biosynthesis, sulfur metabolism and citrate cycle (TCA cycle) (Figure 2A).

**Figure 2.**
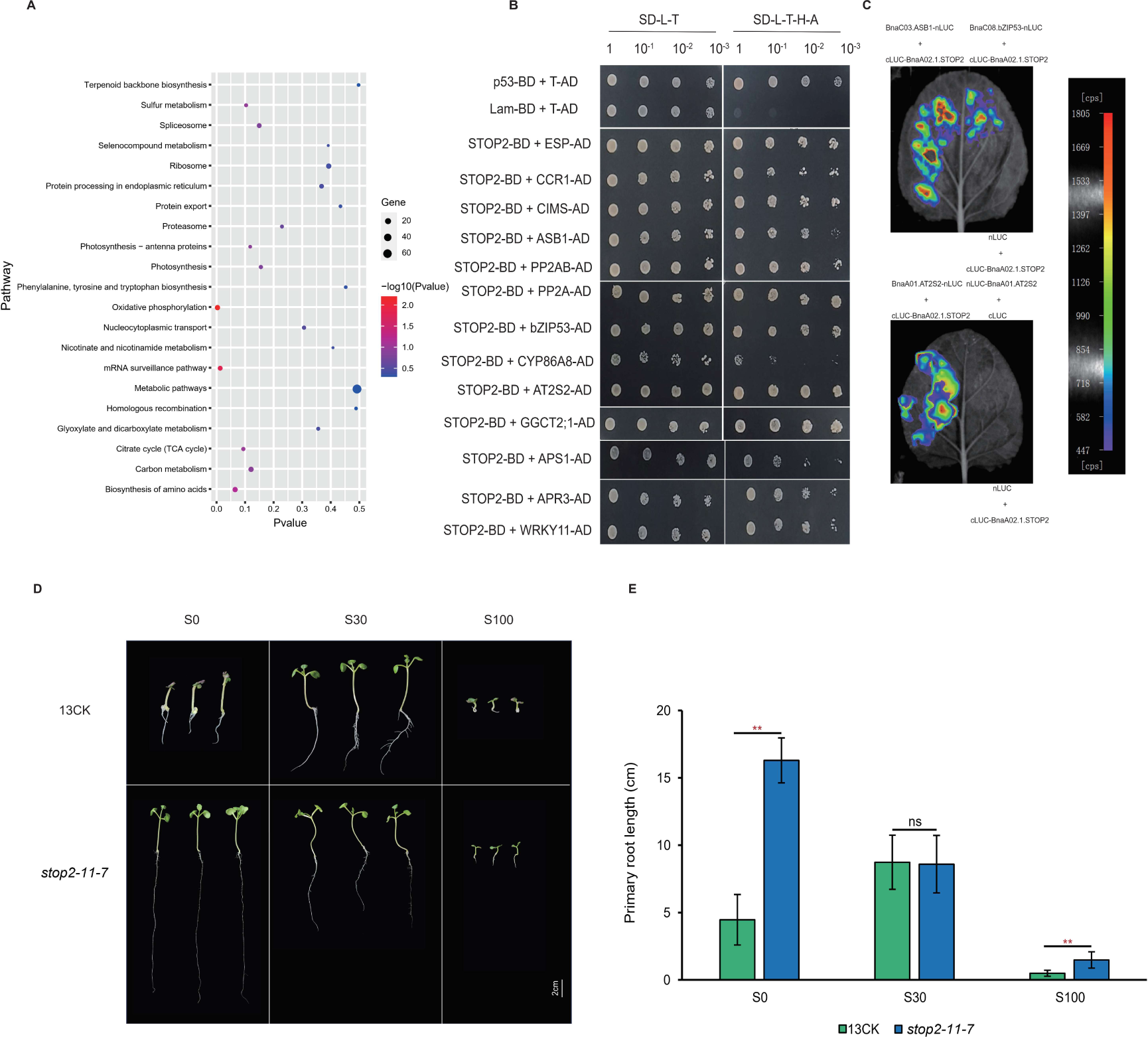
Interaction proteins of BnaA02.1.STOP2 are involved in sulfate metabolism. (**A**) KEGG analysis of interacting proteins with BnaA02.1.STOP2. (**B** and **C**) Screening point-to-point and SLC validation of BnaA02.1.STOP2 Y2H library. (**D** and **E**) Representative 2-week-old WT and *Bnastop2* mutant seedlings grown on media containing increasing concentrations of sulfate (SO^4–^), as indicated. S0, -S medium; S30, including 30 μM S medium; S100, including100 μM S medium. Scale bar represents 2 cm. Differences between mutants and WT are significant at p< 0.05 (*) and p<0.01 (**) by two-tailed Student’s t test (10 ≤ n ≤ 20).

Among these interaction proteins, nine proteins related to sulfur metabolism have been identified more than three times (Figure 2B, Supplemental Figure 4). These genes are distributed across pathways associated with sulfur assimilation, cysteine and methionine biosynthesis, GSL, and glutathione (GSH) metabolism. For example, *BnaA01G0203700ZS* is homologous to Arabidopsis *ATP SULFURYLASE 1* that encodes an ATP sulfurylase, the inaugural catalyst in the sulfate assimilation pathway (Hatzfeld et al., 2000). *BnaA01G0120900ZS* is homologous to Arabidopsis *APS REDUCTASE 3* that encodes a protein disulfide isomerase-like (PDIL) protein and responsible for reducing sulfate for cysteine biosynthesis (Setya et al., 1996). *BnaA04G02*78900ZS is homologous to Arabidopsis *O-ACETYLSERINE (THIOL) LYASE B* that encodes the isoform OASB of O-acetylserine (thiol) lyase (OAS-TL), and facilitates the enzymatic conversion of O-acetylserine and inorganic sulfide into cysteine (Heeg et al., 2008; Wirtz et al., 2010). *BnaA01G0197100ZS* is homologous to Arabidopsis *SERINE/THREONINE PROTEIN PHOSPHATASE 2A* that encodes the B’ regulatory subunit of PP2A (AtB’gamma), involved in S-adenosylmethionine cycle and indole GSL biosynthetic process (Rahikainen et al., 2017). *BnaA05G0156600ZS* is homologous to Arabidopsis *EPITHIOSPECIFIER PROTEIN* that encodes Epithiospecifier protein and involved in GSL catabolic process (Burow et al., 2008; Buxdorf et al., 2013). Moreover, *BnaA09G0061600ZS* is homologous to Arabidopsis *GAMMA-GLUTAMYL CYCLOTRANSFERASE 2;1* (*GGCT2;1*), a gene that regulates GSH metabolism of important sulfur-containing secondary metabolites (Ito et al., 2022; Paulose et al., 2013), and *BnaA01G0165000ZS* is homologous to Arabidopsis *AT2S2*, a catalytic enzyme that regulates sulfur-containing storage proteins (Fujiwara et al., 2002), were also detected in interactions. Interestingly, genes involved in the regulation of plant-pathogen interactions and lipid metabolism pathways were identified. Among them, *BnaA01G0060300ZS* is homologous to Arabidopsis *WRKY11* that participates in defense reactions against bacteria (Ali et al., 2018; Jiang et al., 2016), and *BnaC04G0080200ZS* is homologous to lipid transfer protein *LTP1* (Li-Beisson et al., 2015). Further random verification using split-luciferase complementation (SLC) confirmed the interactions of BnaA08.ASB1, BnaA01.AT2S2, and BnaC08.bZIP53 with BnaA02.1.STOP2 identified in the Y2H screen (Figure 2C). These findings hinted that *BnaSTOP2* encompasses the intricate regulatory network governing sulfur metabolism beyond nitrogen metabolism.

### *BnaSTOP2* is associated with the response sensitivity in both sulfur-deficient and sulfur-excessive conditions

Sulfur metabolism has been unequivocally linked to the proliferation and maturation of roots (Maruyama-Nakashita et al., 2006; Zhao et al., 2014). Then, we observed the root growth of the *Bnastop2* mutants under different levels of sulfur supply for 2 weeks. As shown in Fig. 2D&E, *Bnastop2* seedlings exhibited a three-fold increase in the length of their primary roots and more robust shoot development in comparison to the WT (13CK) without sulfur supply (S0, 0 μM). Under the condition of S30 (30 μM), the primary root length in the WT were comparable to that in *Bnastop2 mutants*; however, when individually compared to that of S0, the WT and *Bnastop2* mutants showed opposite tendency in primary root length, as it has been almost doubled in the WT while notably reduced in *Bnastop2* mutants (Figure 2D-2E). When the sulfur supply reaches a high concentration (S100, 100 μM), the primary root growth of the WT and *Bnastop2* were both severely inhibited; comparatively, *Bnastop2* has a longer primary root than the WT (Figure 2D-2E). Summarized, whether under the sulfur-deficient or sulfur-excessive conditions, the WT were more sensitive to sulfur supply than *Bnastop2* mutants. These findings underscored a crucial role of *BnaSTOP2* in sulfur-mediated primary root growth, implying that *BnaSTOP2* may be a key regulator in plant growth responses to sulfur availability.

### *BnaSTOP2* plays a critical role in maintaining sulfur metabolism in *B. napus*

To comprehensively elucidate the metabolic alterations in roots of *Bnastop2* mutants, we employed a widely targeted metabolomics analysis, quantifying 362 metabolites through LC-MS. Our investigation identified a subset of 36 metabolites exhibiting significant up-regulation and 18 metabolites displaying notable down-regulation (Supplemental Figure 5A-5B, Supplemental Dataset 5). KEGG analysis revealed that these differential metabolites were primarily associated with the biosynthesis of secondary metabolites, GSL biosynthesis, and 2-oxocarboxylic acid metabolism (Figure 3A). A more detailed examination of the differential metabolites unveiled substantial increases in 5-methylthiopentyl GSL, 1,7-Dimethylxanthine, 8-methylthiooctyl GSL, Riboflavin 5’-Adenosine Diphosphate, and L-serine levels, while 3-(Methylthio)propyl GSL levels were significantly reduced in the roots of *Bnastop2* mutants (Supplemental Figure 5C). These findings confirmed the significant impact of *BnaSTOP2* on the overall metabolic profile of roots, with a particularly notable impact on GSL metabolism.

**Figure 3.**
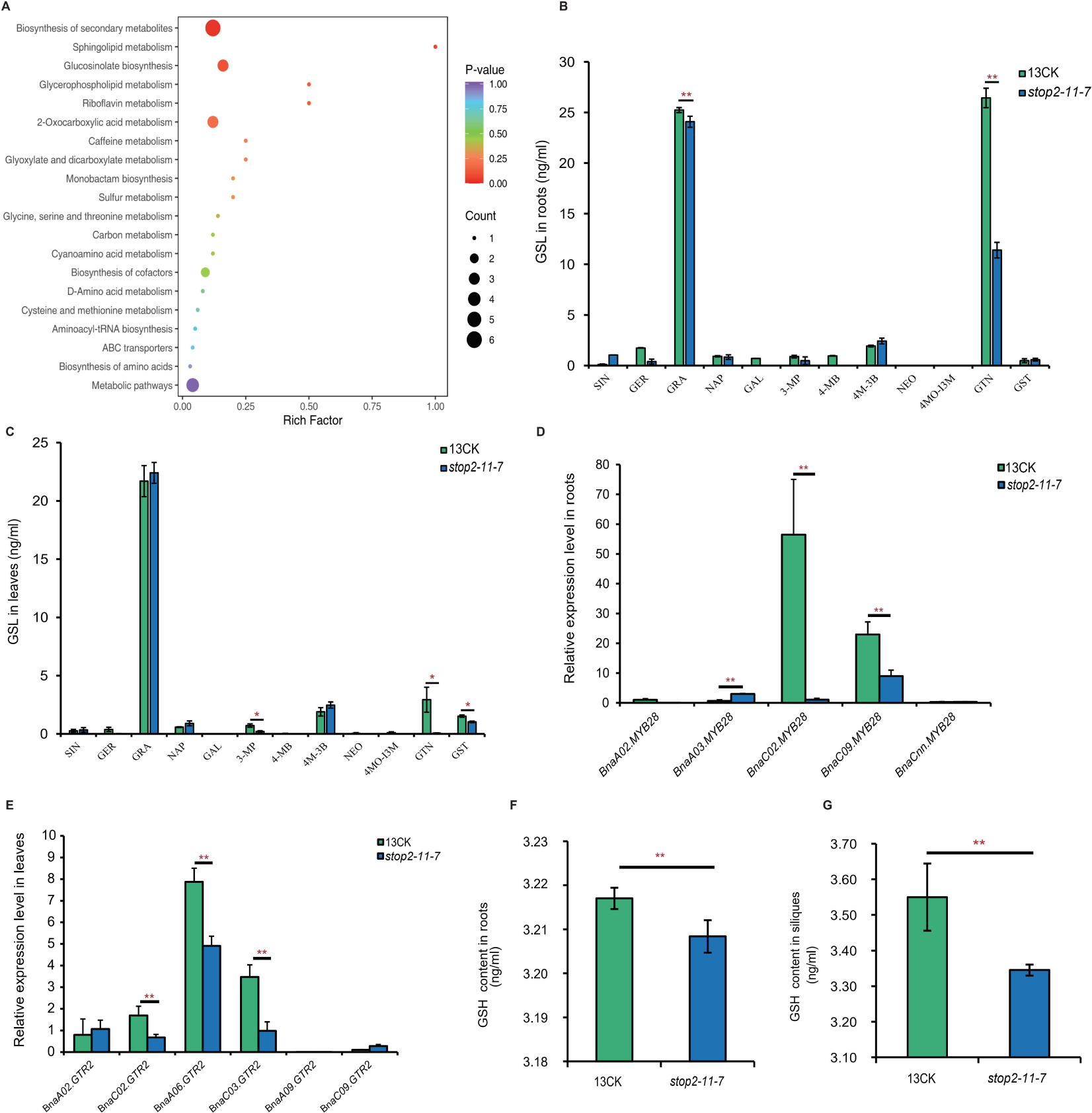
*BnaSTOP2* is important for the accumulation of sulfate metabolites. (**A**) KEGG analysis of differential metabolites in roots of WT and *Bnastop2* mutants. (**B** and **C**) The content of GSL (ng/ml) in roots and leaves of WT and *Bnastop2* mutants. SIN: Sinigrin; GER: Glucoerucin; GRA: Glucoraphanin; NAP: Gluconapin; GAL: Glucoalyssin; 3-MP: Glucoiberverin; 4-MB: Glucosativin; 4M-3B: Glucoraphenin; NEO: Neoglucobrassicin; 4MO-I3M: 4-Methoxyglucobrassicin; GTN: Glucotropaeolin; GST: Gluconasturtiin. (**D** and **E**) Relative RNA level of *BnaMYB28s* and *BnaGTR2s* in WT and *Bnastop2* mutants. (**F** and **G**) Comparisons of GSH (ng/ml) in roots and siliques of WT and *Bnastop2* mutants. For (**B**) to (**G**), differences between mutants and WT are significant at p< 0.05 (*) and p<0.01 (**) by two-tailed Student’s t test.

Sulfur-containing secondary metabolites, such as GSL and GSH, are integral components of the sulfur reservoir in plants (Jez, 2019; Venditti and Bianco, 2020). Thus, we measured the GSL content in various tissues of the *Bnastop2* mutants by LC-MS. We quantified a total of 12 GSLs, encompassing aliphatic GSLs (SIN-Sinigrin, GER-Glucoerucin, GRA-Glucoraphanin, NAP-Gluconapin, GAL-Glucoalyssin, 3-MP-Glucoiberverin, 4-MB-Glucosativin, 4M-3B-Glucoraphenin), indolic GSLs (NEO-Neoglucobrassicin, 4MO-I3M-4-Methoxyglucobrassicin), and aromatic GSLs (GTN-Glucotropaeolin and GST-Gluconasturtiin) (Supplemental Figure 6, Supplemental Dataset 6). Among the roots of *Bnastop2* mutants, compared to WT (13CK), only the content of GRA and GTN displayed a significant decrease (Figure 3B). In the leaves, there was a notable reduction observed in the content of aromatic GSL (GTN and GST) (Figure 3C). In addition, the seed protein content of *Bnastop2* mutants also significantly decreased (Supplemental Figure 7). We also quantified the GSH content in *Bnastop2* mutants by LC-MS (Supplemental Figure 8A, Supplemental Dataset 7). Compared to WT (13CK), a significant reduction in GSH levels were observed in the roots and siliques of the *Bnastop2* mutants (Figure 3F-3G). However, no substantial difference was observed in the leaves (Supplemental Figure 8B). These findings indicated that the mutation of *BnaSTOP2s* altered the content of GSL and GSH content across various tissues.

Besides, considering that *BnaMYB28s* and *BnaGTR2s* (*GLUCOSINOLATE TRANSPORTER-2*) are crucial for GSL accumulation in *B. napus* (Tan et al., 2021; Zhou et al., 2023), we analyzed their expression levels in the roots and leaves of *Bnastop2* mutants by RT-qPCR. In the roots, the expression levels of *BnaA02.GTR2, BnaC02.MYB28* and *BnaC09.MYB28* were significantly down-regulated, while the expression of *BnaA03.MYB28* was significantly up-regulated; in the leaves, the expression of *BnaC09.MYB28*, *BnaC02.GTR2*, *BnaA06.GTR2* and *BnaC03.GTR2* were significantly down-regulated in leaves (Figure 3D-3E, Supplemental Figure 9, Supplemental Dataset 8). These results suggested that *BnaSTOP2* may be a core factor in maintaining sulfur metabolism in *B. napus*.

### Knockout of *BnaSTOP2s* greatly changes the expression profiles of multiple sulfur metabolism genes in both source and sink tissues

The 20-day period after flowering (20 DAF) is a critical phase of reproductive growth in *B. napus*, characterized by intensive nutrient absorption and energy metabolism (Yu et al., 2010). Therefore, we conducted a comprehensive transcriptome analysis focusing on both WT (13CK) plants and *Bnastop2* mutants across roots and siliques at 20 DAF. These tissues serve as pivotal sources and sinks for sulfur metabolism, respectively (Hawkesford, 2012; Miller and Chapman, 2011). In our investigation of roots, we identified 3067 differentially up-regulated genes and 3378 differentially down-regulated genes (|log_2_foldchange | ≥2 and pval ≤0.05) (Supplemental Figure 10A). Subsequent KEGG enrichment analysis showing that both up-regulated and down-regulated genes were mainly enriched in sulfur metabolism, nitrogen metabolism, and fatty acids metabolism (Figure 4A, Supplemental Figure 10B, Supplemental Dataset 10 and 11). Within the siliques, a total of 7,726 DEGs were identified (|log2foldchange | ≥2 and pval ≤0.05) (Supplemental Figure 11A). The KEGG analysis of DEGs revealed their predominant enrichment in pathways such as cysteine and methionine metabolism, phenylalanine, and tryptophan biosynthesis, as well as sulfur metabolism (Figure 4F, Supplemental Figure 11B and 11C, Supplemental Dataset 12). These findings suggested that the mutation of *BnaSTOP2* had a substantial impact on the metabolic processes occurring in the roots and siliques.

**Figure 4.**
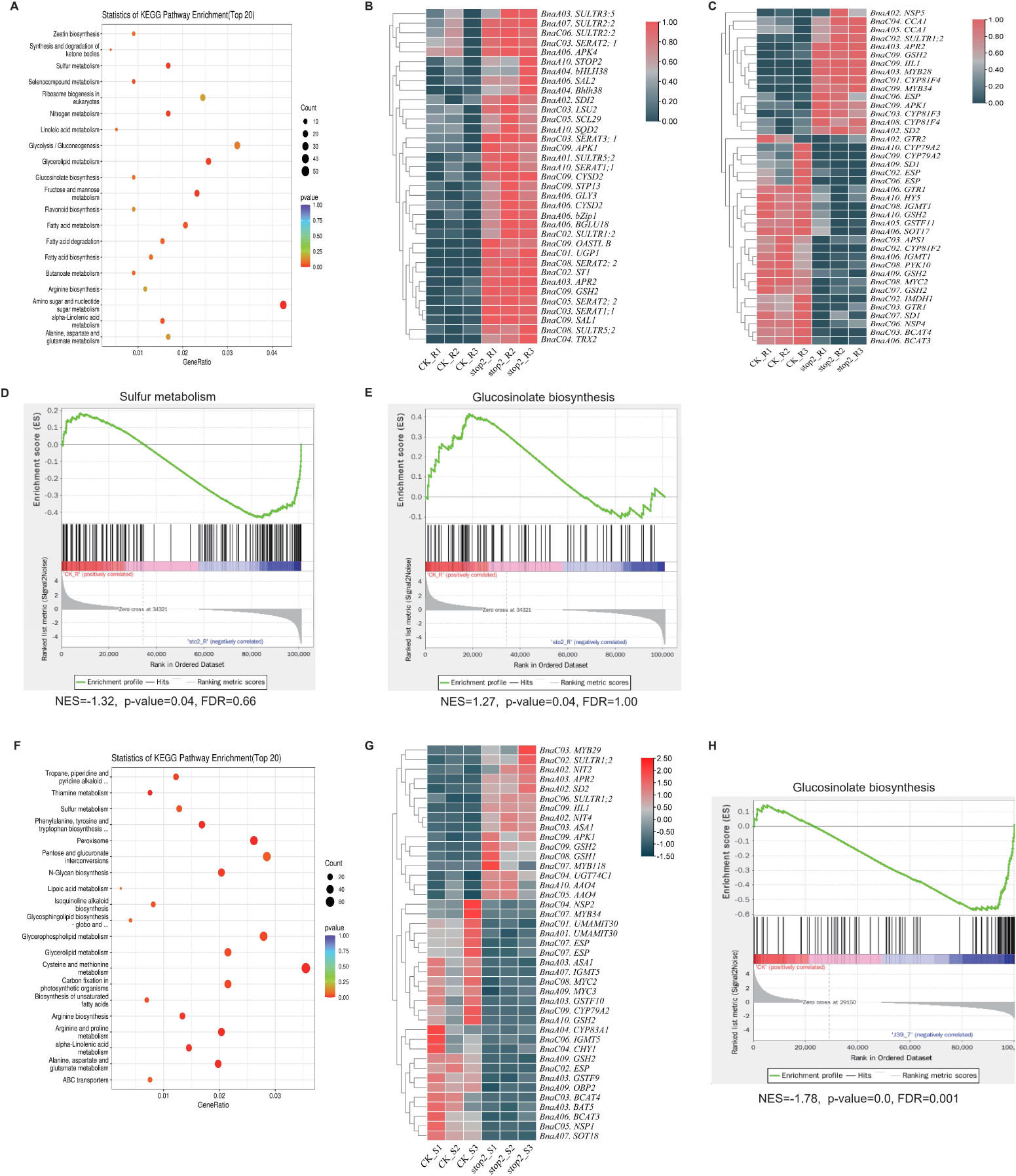
*Bnastop2* mutants DEGs are significantly enriched in amino acid and sulfur metabolism pathway. (**A**) KEGG analysis of down-regulated genes in roots of *Bnastop2* mutants and WT. (**B**) Up-regulated genes of sulfur metabolism related genes in roots of *Bnastop2* mutants and WT. (**C**) DEGs of GSL metabolism related genes in roots of *Bnastop2* mutants and WT. (**D** and **E**) GSEA analysis of the KEGG terms enriched in up- or down-regulated genes in the roots of *Bnastop2* mutants compared with the WT. NES, p-value and FDR value are presented. (**F**) KEGG analysis of differentially expressed genes in siliques of *Bnastop2* mutants and WT. (**G**) DEGs of GSL metabolism related genes in siliques of *Bnastop2* mutants and WT. (**H**) GSEA analysis of the KEGG terms enriched in up- or down-regulated genes the siliques of *Bnastop2* mutants compared with the WT. GSEA, Gene Set Enrichment Analysis; NES, normalized enrichment score.

In roots of *Bnastop2* mutants, our findings showed that 35 genes associated with sulfur metabolism exhibited significant up-regulation, while 25 genes related to sulfur metabolism showed significant down-regulation (Figure 4B, Supplemental Figure 10C, Supplemental Dataset 13). Moreover, several genes belonging to the sulfate transporter family exhibited up-regulation. On the other hand, previous studies have emphasized the significance of sulfur metabolism in orchestrating the complex mechanisms linked to GSL metabolism (Aarabi et al., 2020; Zhang et al., 2020). Notably, the up-regulated genes included important GSL-associated genes, such as *BnaA03.MYB28* and *BnaC09.MYB34*, whereas transporters like *BnaA06.GTR1 (GLUCOSINOLATE TRANSPORTER-1) and BnaA02.GTR2* were predominantly down-regulated in root of *Bnastop2* mutants (Figure 4C, Supplemental Dataset 14). In addition, in siliques, we observed a notable up-regulation of 16 genes implicated in GSL metabolism, along with a down-regulation of 25 genes linked to GSL metabolism. Key regulatory genes that play crucial roles in both GSL and sulfur metabolism pathways, such as *BnaA03.APR2*, *BnaC08.GSH1* (*GLUTATHIONE SYNTHETASE 1*)*, BnaC09.GSH2* (*GLUTATHIONE SYNTHETASE 2*), *BnaC09.APK1* (*APS kinase 1*), and *BnaSULTR1;2s*, exhibited significant up-regulation. Notably, we also observed a substantial down-regulation of pivotal regulatory genes implicated in the biosynthesis of aliphatic GSL, including *BnaC07.MYB34*, *BnaC08.MYC2*, *BnaA09.MYC3*, *BnaA06.BCAT3 (BRANCHED-CHAIN AMINOTRANSFERASE 3)*, *BnaA03.BCAT4* (*BRANCHED-CHAIN AMINOTRANSFERASE 4*), and *BnaA03.BAT5* (*BILE ACID TRANSPORTER 5*) (Figure 4G, Supplemental Dataset 15).

Furthermore, to parse the overall changes in sulfur metabolism-related pathways, we performed Gene Set Enrichment Analysis (GSEA) to identify gene sets with synergistic differences (Subramanian et al., 2005). In the roots of *Bnastop2* mutants, we observed a down-regulation trend in sulfur metabolism, and an up-regulation trend in GSL biosynthesis (Figure 4D-4E). Furthermore, we observed a down-regulation trend in the expression of the GSL metabolism-related gene set in siliques (Figure 4H). However, it is worth noting that crucial genes implicated in the biosynthesis and transportation of GSL were found to be down-regulated, specifically *BnaA01.UMAMIT30* and *BnaC01.UMAMIT30* (Figure 4G), which shares homology with Arabidopsis *USUALLY MULTIPLE ACIDS MOVE IN AND OUT TRANSPORTERS 30* (*UMAMIT30*), exhibits significantly decreased expression levels. *UMAMIT30* is recognized for its expression in the funicle of reproductive tissue and its indispensable contribution to the translocation of GSL into seeds in Arabidopsis (Xu et al., 2023). Taken collectively, these findings suggested that *BnaSTOP2* exerts significant regulatory influences on diverse facets of sulfur metabolism, encompassing sulfate transport, sulfate assimilation, GSL biosynthesis, as well as cysteine and methionine metabolism at both source and sink.

### *BnaSTOP2* positively regulates *S. sclerotiorum* resistance in *B. napus*

Essential sulfur-containing primary metabolites, secondary metabolites, and proteins play a critical role in plant defense against pathogens, directly or indirectly bolstering defense responses (Venditti and Bianco, 2020). Through Y2H analysis, we provided evidence supporting the critical involvement of *BnaSTOP2* in defense regulation, including pathogen resistance (Figure 2B). As a necrotrophic fungal pathogen, *S. sclerotiorum* causes the most severe yield loss in many rapeseed grown regions(Kamal et al., 2016). To analyze the effects of *BnaSTOP2* on defense response in *B. napus*, *S. sclerotiorum* was inoculated onto the leaves of *Bnastop2* mutants. A significant increase in lesion size was observed in the mutant at 48 hours post-inoculation (hpi) (Figure 5A and 5B, Supplemental Figure12). Additionally, we generated *BnaA02.1.STOP2 OE* (*BnaSTOP2-OE*) mutants in Xiaoyun, which is susceptible to *S. sclerotiorum*. The expression level of *BnaA02.1.STOP2* significantly increased in OE mutants (Supplemental Figure 13). In comparison to the WT, *BnaSTOP2-OE* mutants demonstrated enhanced resistance (Figure 5C and 5D). These results indicated that *BnaSTOP2s* may positively regulate the resistance to *S. sclerotiorum* in *B. napus*.

**Figure 5.**
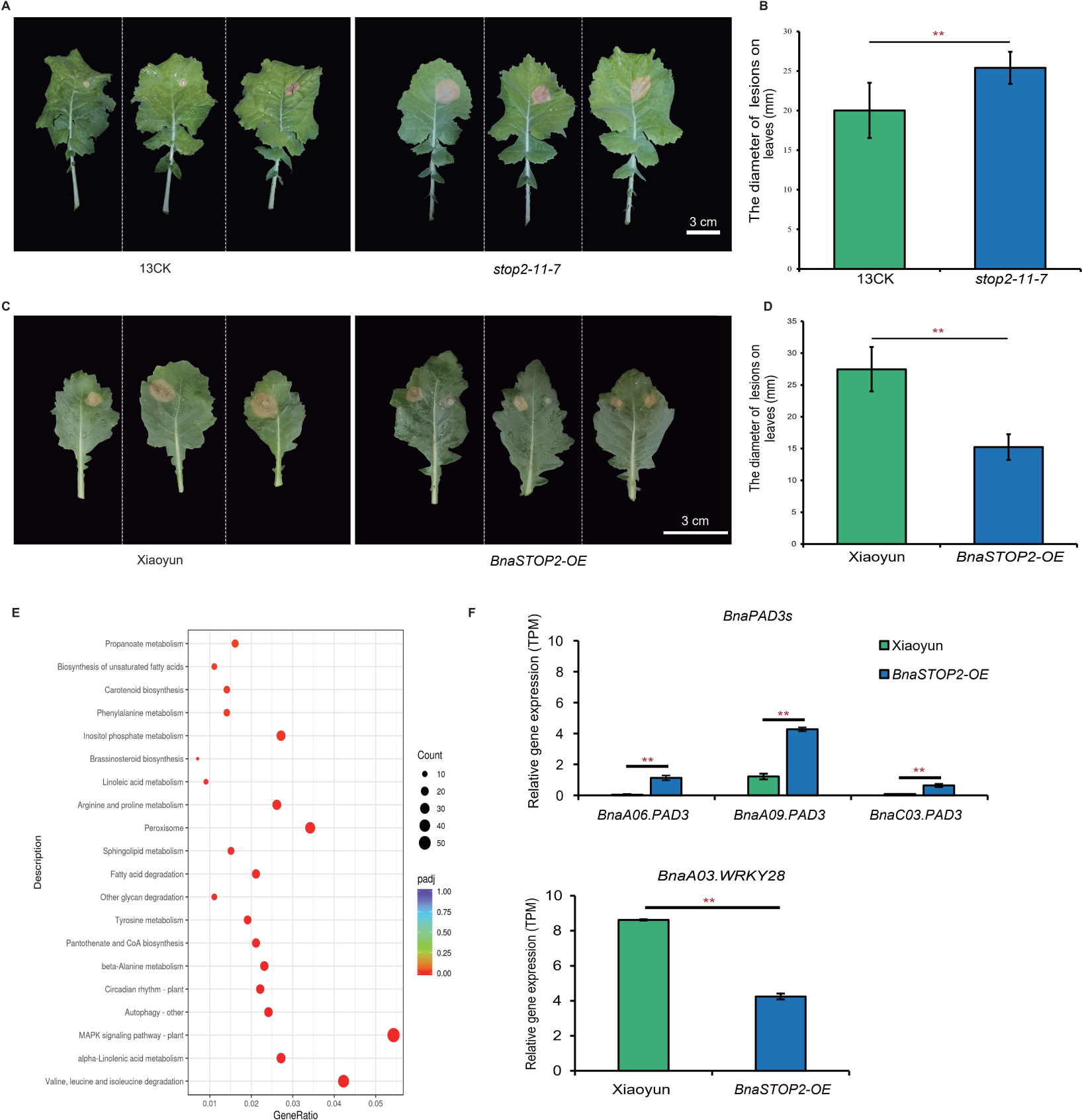
*BnaSTOP2* has an important impact on defense response. (**A** and **B**) Identification of resistance to *Sclerotinia sclerotiorum*, on leaves of *Bnastop2* mutants (n=10). (**C** and **D**) Identification of resistance to *Sclerotinia sclerotiorum*, on leaves of *BnaSTOP2-OE* mutants (n=14). For (**B)** and (**D**), differences between mutants and WT are significant at p< 0.05 (∗) and p<0.01 (∗∗) by two-tailed Student’s t test. (**E**) KEGG analysis of up-regulated genes in *BnaSTOP2-OE* mutants and Xiaoyun (48 hpi). (**F**) Normalized gene expressed levels (TPM) from RNAseq data for *BnaPAD3s* and *BnaA03.WRKY28* in *BnaSTOP2-OE* mutants and Xiaoyun (48 hpi). Differences between mutants and Xiaoyun are significant at p< 0.05 (∗) and p<0.01 (∗∗) by two-tailed Student’s t test.

Furthermore, RNA-seq analyses were performed at 0 hpi and 48 hpi respectively. Subsequent analysis of KEGG revealed that, up-regulated genes in *BnaSTOP2-OE* mutants at 48 hpi and 0 hpi, primarily enriched in the MAPK signaling pathway, cysteine and methionine metabolism, GSH metabolism, sulfur metabolism, and GSL biosynthesis (Supplemental Figure 14A, Supplemental Dataset 16); and up-regulated genes between *BnaSTOP2-OE* mutants (48 hpi) and Xiaoyun (48 hpi) mainly enriched in MAPK signaling pathway (Figure 5E). Besides, the GSEA analysis of *BnaSTOP2-OE* (48 hpi) revealed an ascending pattern in the metabolic pathways of cysteine and methionine, ABC transporters, and the biosynthesis of diverse secondary metabolites in plants (Supplemental Figure 15B-15D). Moreover, in *BnaSTOP2-OE* mutants at 48 hpi, several crucial genes associated with *S. sclerotiorum* resistance exhibited differential expression. For instance, *PHYTOALEXIN DEFICIENT 3* (*PAD3*), which encodes a key enzyme involved in camalexin biosynthesis, is known to inhibit the growth of *S. sclerotiorum* (Purnamasari et al., 2015). In *BnaSTOP2-OE* mutants at 48 hpi, *BnaPAD3* (A6/A9/C3) displayed significant up-regulation. Conversely, *BnaA03.WRKY28*, a negative regulator of *S. sclerotiorum* resistance in *B. napus* (Zhang et al., 2022), exhibited significant down-regulation (Figure 5F). These findings substantiated the proposition that *BnaSTOP2* plays a constructive role in enhancing *S. sclerotiorum* resistance in *B. napus*. The enhanced resistance observed in *BnaSTOP2*-OE mutants can be attributed to the upregulation of genes associated with the MAPK signaling pathway, cysteine and methionine metabolism, GSH metabolism, sulfur metabolism, and GSL biosynthesis. Furthermore, crucial genes involved in camalexin production and negative regulation of *S. sclerotiorum* resistance further supported the role of *BnaSTOP2s* in plant defense mechanisms.

## DISCUSSION

The C2H2 zinc finger transcription factor family includes the pivotal member *STOP2*, which in Arabidopsis, displays a unique homologous, *STOP1*. This study extends this understanding by uncovering the emerging role of *STOP2* in modulating sulfur metabolism within *B. napus*. Inorganic sulfate entry into plants typically involves absorption, reduction to sulfide, and subsequent integration into crucial molecules like cysteine, methionine, GSH, and GSL (Brosnan and Brosnan, 2006; Mugford et al., 2012). To investigate the impact of *BnaSTOP2s* on sulfur metabolism, we analyzed the levels of GSH and GSL in the *Bnastop2* mutants. It was observed that the roots and siliques of the mutants showed a significant decrease in GSH content. Additionally, the *Bnastop2* mutants displayed significant changes in the levels of various metabolites in the roots, with noticeable variations in the levels of different GSL in both the roots and leaves (Figure 3B-3C). Furthermore, *BnaMYB28s* and *BnaGTR2s* constitute pivotal determinants for GSL accumulation in *B. napus*, and their expression levels both significantly decreased in *Bnastop2* mutants (Figure 3D-3E). Therefore, it is speculated that *BnaSTOP2* potentially modulates accumulation through the regulation of *BnaMYB28s* and *BnaGTR2s*.

Moreover, the primary root length of the *Bnastop2* mutants grew 3.82-fold longer than WT on sulfur deficiency medium (Figure 2D-2E). This is consistent with the change in root length of *ggct2;1* under sulfur deficient conditions (Joshi et al., 2019), and BnaC09G0050100ZS (homologous to GGCT2;1) is interact with BnaA02.STOP2 (Figure 2B), thus implying that *BnaSTOP2s* may participate in the changes in root structure during sulfur deficiency reactions together with GGCT2;1. Furthermore, under high sulfur conditions, the primary root length of the *Bnastop2* mutants remains notably longer than that of the WT. Consequently, we speculated that *BnaSTOP2* controls the dynamic balance of sulfur metabolism. This regulation ensures a stable sulfur metabolism across diverse environmental conditions, diminishing sulfur content under high sulfur conditions and elevating it under low sulfur conditions.

As plants rely on sulfate uptake from the soil through SULTRs, sulfate is subsequently distributed within the plant cell, forming a dynamic "sulfur pool". The metabolic process converts sulfate into diverse primary and secondary metabolites. The sulfur assimilation pathway is integral to the synthesis of sulfur-containing amino acids and secondary metabolites, including GSL (Hawkesford and De Kok, 2006). The modulation of gene expression of ATP sulfurylase (ATPS), APS reductase (APR), and APS kinase (APK) facilitates the biochemical allocation of sulfur across various branches of the assimilation pathway (Mugford et al., 2009; Vauclare et al., 2002). In the roots of *Bnastop2* mutants, DEGs are significantly enriched in sulfur metabolism pathway, for instance, the expression of the sulfur metabolism gene set is generally down-regulated (Figure 4A and 4D). Genes involved in sulfur metabolism such as *ATPS1*, *APR52*, *APK3*, and *GSH2* all showed down-regulated. However, sulfate transporters exhibit up-regulated expression, such as *SULTR1; 1*, *SULTR2; 2*, *SULTR3; 5*, *SULTR5; 2* (Figure 4C), perhaps owing to the mutation of *BnaSTOP2*, the sulfur metabolism in roots is disrupted, and as roots serve as a primary sulfur source, can direct influence plant growth. To alleviate this situation, plants enhance the expression of sulfur transporters, facilitating the transportation of sulfate from the soil to replenish the sulfur source.

On the other hand, siliques have known as a vital site for photosynthesis during seed maturation and a significant sink for storing synthesized carbohydrates from vegetative tissues (Samizadeh et al., 2007). In the siliques of *Bnastop2* mutants, the key regulatory genes involved in GSL biosynthesis and transportation displayed a significant down-regulation. These genes include *BnaC07.MYB34*, *BnaC08.MYC2*, *BnaA09.MYC3*, *BnaA06.BCAT3*, *BnaA03.BCAT4*, *BnaA03.BAT5*, *BnaA01.UMAMIT30*, and *BnaC01.UMAMIT30* (Figure 4G). Therefore, there was a downward trend in the expression of the GSL metabolism gene set in the siliques of *Bnastop2* mutants. In addition, in combination with Y2H, we screened regulatory factors at different stage of sulfur metabolism that interact with BnaA02.1.STOP2 (Figure 1C-1D). Our investigation into the role of *BnaSTOP2* in sulfur metabolism has uncovered its significant participation in both source and sink tissues.

Sulfur metabolism is intricately involved in a variety of defense reactions. Cysteine, a precursor of sulfur defense compounds, holds a central position in these reactions (Chan et al., 2019). Secondary metabolites containing sulfur, such as GSL, have been identified as potent contributors to resistance against pathogens like *S. sclerotiorum*. Our experimental data supported this, revealing a significant decrease in resistance of *Bnastop2* mutants upon inoculation with *S. sclerotiorum* (Figure 5A-5B), as opposed to the enhanced resistance observed in *BnaSTOP2* overexpressing mutants compared to the WT (Xiaoyun) (Figure 5C-5D). From the substantial decrease in Glucoiberverin (3-MP) and Gluconastutin (GST) content within the leaves of the *Bnastop2* mutant, along with the positive correlation between these two GSLs and resistance against *S. sclerotiorum* (Abuyusuf et al., 2018; Teng et al., 2021). This led us to propose the potential involvement of *BnaSTOP2* in regulating *S. sclerotiorum* resistance through the modulation of 3-MP and GST. On the other hand, our analysis revealed a substantial enrichment of the MAPK immune signaling pathway at 48 hpi (Figure 5E). This observation is complemented by the significant up-regulation of *BnaPAD3* (*A6/A9/C3*) and the noteworthy down-regulation of *BnaA03.WRKY28* in *BnaSTOP2-OE* mutants at 48 hpi (Figure 5F). These findings introduced a dynamic layer to our understanding, hinting at an interplay between *BnaSTOP2* and the complex network of signaling pathways crucial for plant immunity.

In summary, our study suggests that *BnaSTOP2* plays a crucial role in regulating sulfur metabolism and defense responses. It influences sulfur assimilation levels, sulfur compound metabolism in roots, leaves, and siliques, and pathogen resistance in leaves by forming multiple protein complexes (Figure 6). This newly discovered regulator has shown that it can modulate sulfur assimilation and metabolism to improve pathogen resistance, offering essential information for developing breeding strategies aimed at producing high-quality and high-yield *Brassica* crops.

**Figure 6.**
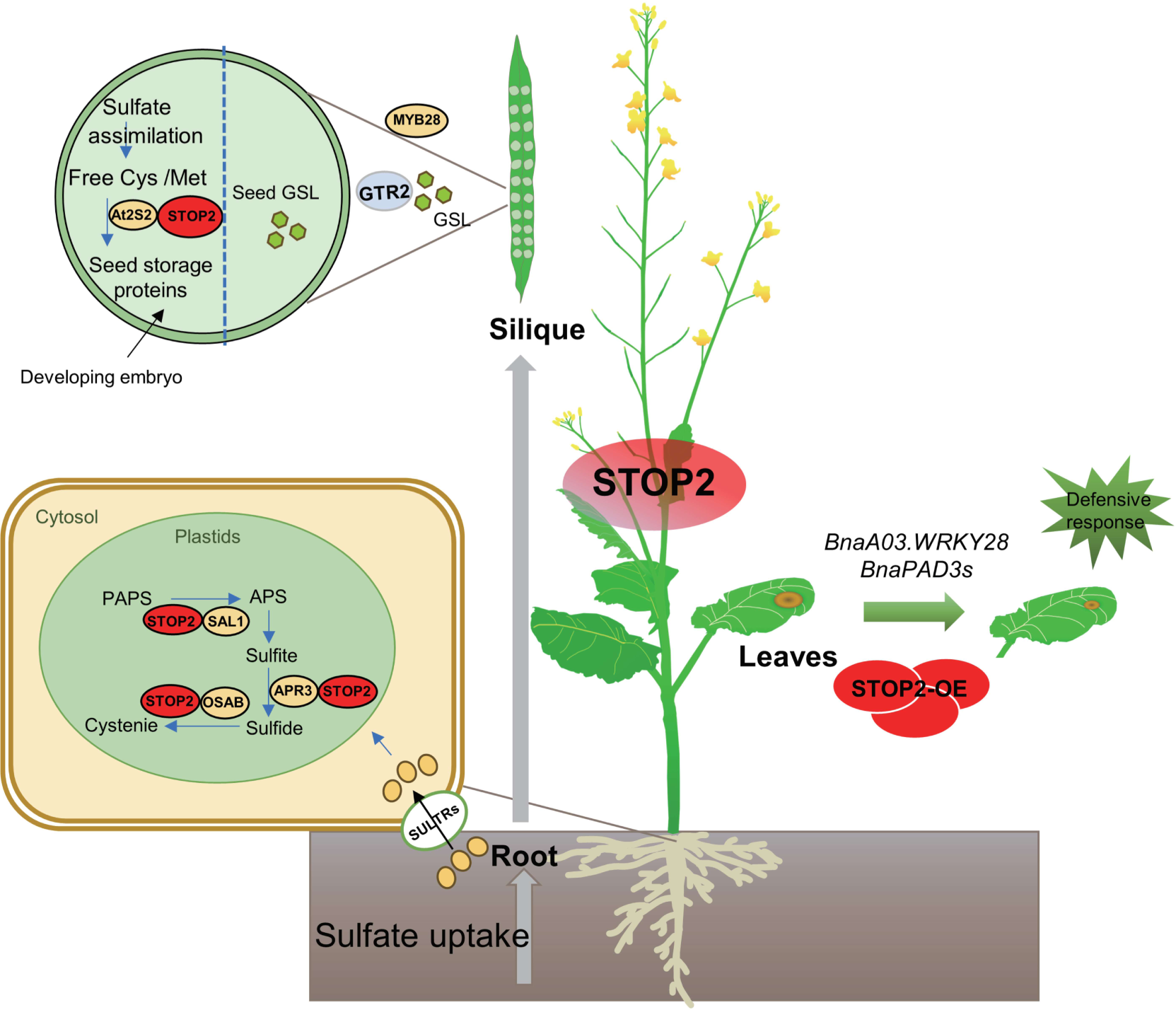
Zinc finger transcription factor *STOP2* is a new regulator of sulfur metabolism and *Sclerotinia sclerotiorum* resistance in *Brassica napus.* During the process of sulfur absorption in the root system, *BnaSTOP2* affects sulfur absorption by affecting the expression levels of sulfur transporters (SULTRs), as well as sulfur metabolism in sulfur roots by affecting the expression levels of *SAL1*, *OASB*, and *APR3*. Moreover, when the siliques undergo sulfur pooling, *BnaSTOP2* exerts influence over the accumulation of GSL by modulating the expression of *BnaMYB28s* and *BnaGTR2s*. *BnaSTOP2* also impacts the accrual of stored proteins in seeds through interactions with At2S2.Additionally, in the face of *S. sclerotiorum* -induced leaf damage, *BnaSTOP2* enhances the plant’s resistance to diseases by effectively modulating the expression of *BnaPAD3s* and *BnaA03.WRKY28*.

## MATERIALS and METHODS

### Plant materials

Mutants were derived from *B. napus* varieties named 13CK or Xiaoyun. Overexpression (OE) and CRISPR/Cas9 lines were established, including *Bnastop2*/13CK, *BnaA02.1.STOP2-OE*/ Xiaoyun. Mature open-pollinated seeds from the *Bnastop2* mutants and WT (13CK) were collected and desiccated to analysis of seed GSL content and seed protein content. The analysis was conducted utilizing a Foss NIRSystems 5000 spectroscope, which operates based on the principle of near-infrared reflectance (Gan et al., 2003). Each accession underwent analysis with five replications. The measurements were carried out at the esteemed National Research Center of Rapeseed Engineering and Technology, located within the prestigious grounds of Huazhong Agricultural University in the bustling city of Wuhan, China.

### Construction of vector and plant transformation

The CRISPR/Cas9 genome editing system, employed in this study to edit the *BnaSTOP2* genes, was established based on established procedures (Xing et al., 2014). The primers utilized for the assembly of the sgRNA vector are documented in Supplementary Table S17. The OE construct was driven by CaMV35S (pCAMBIA2300 vector), the primers used for the construction of the OE vector are listed in the Supplemental Table S17. In this study, the donor plants for CRISPR/Cas9 and overexpression (OE) techniques were the ’13CK’ variety, characterized by its wild-type properties as a semi-winter-type rapeseed with a seed GSL content of 137.25 μmol/g and an erucic acid composition of 34.26% in the seeds, and the ’Xiaoyun’ variety, also a wild-type but spring-type rapeseed with a seed GSL content of 22.68 μmol/g and an erucic acid composition of 3.69% in the seeds. The T_0_ transgenic plants were cultivated within a greenhouse, wherein a light and dark cycle of 16 and 8 hours respectively, was maintained at an ambient temperature of 23-25°C. Subsequently, the T_2_ *Bnastop2* mutants, alongside the WT plants, were cultivated in the transgenic experimental field of Huazhong Agricultural University. The leaves of both the WT and mutant plants were harvested for DNA extraction, employing the Transzol sampling method.

### RNA extraction and RT-qPCR

Total RNA was isolated from 100 mg of plant tissue using the plant RNA extraction kit. (Promega, LS1040, China). The RNA-seq samples consisted of *Bnastop2* mutants and WT plants, each with three biological replicates. A quantity of two micrograms of total RNA was employed for cDNA synthesis, accomplished through the utilization of the HiScript 1st Strand cDNA Synthesis Kit (Vazyme, R111-01, China). The RT-qPCR analysis was carried out utilizing the CFX96 Real-Time system (Bio-Rad, USA), employing the ChamQ Universal SYBR qPCR Master Mix (Vazyme, Q711, China). The 2^(-ΔΔCt) method, also known as the ΔCt method, was utilized to compute the relative expression levels of genes. This was performed using three biological samples and three technical replicates per sample (Livak and Schmittgen, 2001). The RT-qPCR primers are provided in Supplemental Table S17. All values were standardized to the transcript levels of the ACTIN7 gene (AT3G46520).

### Y2H

Y2H assays were conducted following the protocols outlined in the Matchmaker GAL4 Two Hybrid System (Clontech). Clone the full-length CDS of each target gene into pGADT7 and pGBKT7 vectors, transfer the non-toxic and non-self-activation bait vector and the recombinant plasmid connected to the pGADT7 vector into AH109 strain according to the yeast transformation method. Y2H assays were conducted in accordance with the Matchmaker GAL4 Two Hybrid System. Clones were cultured on selective dropout medium, either lacking Trp and Leu or Trp, Leu, His, and Ade. The positive controls consisted of pGBKT7-53 and pGADT7-T, while the negative controls included pGBKT7-lam and pGADT7-T. The primer details can be found in Supplemental Table S17.

### SLC

The coding sequence fragments of each target gene were amplified and inserted into the JW771-nLUC and JW772-cLUC vectors. The bacterial suspensions were prepared and delivered into *N. benthamiana* leaves using a needleless syringe, following a methodology established in a prior investigation (Chen et al., 2008). The luminescent signal was captured utilizing an imaging apparatus (NightShade LB 985, Berthold). The primer details can be found in Supplemental Table S17.

### Metabolomics analysis

The procedure for sample preparation and extraction was carried out as follows: Biological specimens underwent cryodesiccation utilizing a Scientz-100F vacuum freeze-dryer. Subsequently, the desiccated sample was fragmented using a Retsch mixer mill (MM 400) equipped with a zirconia bead, operating at a frequency of 30 Hz for a duration of 1.5 minutes. A quantity of 100 mg of the freeze-dried powder was dissolved in 1.2 ml of a 70% methanol solution. The resultant mixture underwent agitation for 30 seconds at 30-minute intervals, repeated a total of six times. Afterward, the sample was stored overnight in a refrigerator at a temperature of 4°C. Following centrifugation at 12,000 rpm for 10 minutes, the extracts were filtered through an ANPEL SCAA-104 filtration system, prior to UPLC-MS/MS analysis. The analytical parameters were ascertained as follows: The UPLC column employed was the Waters ACQUITY UPLC HSS T3 C18 (1.8 µm, 2.1 mm*100 mm) with a column temperature of 40°C. The flow rate was upheld at 0.4 mL/min, and an injection volume of 2 μL was utilized. The solvent system consisted of an aqueous solution with 0.1% formic acid and an acetonitrile solution with 0.1% formic acid. The gradient program that was implemented followed this pattern: 95:5 volume/volume (V/V) at 0 minutes, 5:95 V/V at 10.0 minutes, 5:95 V/V at 11.0 minutes, 95:5 V/V at 11.1 minutes, and 95:5 V/V at 15.0 minutes. The acquisition of scans was performed using a QTRAP triple quadrupole-linear ion trap mass spectrometer, utilizing both the Linear Ion Trap (LIT) and triple quadrupole (QQQ) modes. The instrument was equipped with an ESI Turbo Ion-Spray interface and operated in both positive and negative ionization modes. The QQQ scans were conducted as multiple reaction monitoring (MRM) experiments, with the collision gas (nitrogen) set at a medium level. The selection of specific MRM transitions was further refined through adjustments in the differential pressure (DP) and collision energy (CE).

### Analysis of GSL and GSH using LC–MS/MS

The levels of GSL in the roots, leaves, and seeds of the WT (13CK) and *Bnastop2* mutants were scrutinized. The investigation involved the application of three biological replicas. Subsequent to prompt freezing using liquid nitrogen, the seeds, freshly roots and leaves were preserved at an extreme temperature of -80°C until the extraction of the GSL compounds was carried out. The procedure for GSL extraction strictly adhered to the pre-established protocol (Tan et al., 2021). 2-propenyl GSL (Sinigrin, Sigma-Aldrich) was utilized as an external standard. The analysis of GSL was conducted using LC–MS sec6500 plus Qtrap (Sciex, USA) by following the methods described in previous studies (Nour-Eldin et al., 2012). Separation was achieved on the ACQUITY UPLC BEH C18 Column (130Å, 1.7 µm, 2.1 mm X 100 mm, 1/pk). A solution of formic acid with a concentration of 0.1% (v/v) in water, along with acetonitrile containing 0.1% (v/v) formic acid, served as mobile phases A and B, respectively. The elution profile was as follows: 0–0.5 minutes, 2% B; 0.5 to 7.5 minutes, 2–40% B; 7.5 to 8.5 minutes, 40–90% B; 8.5 to 11.5 minutes, 90% B; 11.6 to 15.5 minutes, 2% B. The flow rate of the mobile phase was set at 300 µL/min. The voltage applied to the ion spray was held at 4,000 V while operating in the negative-ion mode. The temperature of the column was maintained at 40°C, and active exhaust was consistently engaged. Monitoring of analyte precursor ion to fragment ion transitions was carried out using multiple reaction monitoring (MRM). Specific mass transition values can be found in Supplemental Figure 6.

Extraction of reduced GSH: The procedure for extracting reduced glutathione involved preheating 2.0g of the sample at 75 ℃. Subsequently, a 70% methanol-water solution, also preheated to 75 ℃, was added to the sample. The mixture was then subjected to extraction in a 75 ℃ water bath for a duration of 10-30 minutes. During the extraction process, shaking was performed. Following extraction, the mixture was centrifuged to collect the supernatant. This procedure was repeated 2-3 times to ensure efficient precipitation, and the collected supernatants were combined. The extraction solution was further subjected to adsorption treatment using DEAE Sephadex A-25 resin to obtain a comprehensive sample. Following the completion of the filtration process, the sample underwent membrane filtration and was subsequently stored for future utilization. The task of separation was accomplished by employing the ACQUITY UPLC BEH C18 Column, distinguished by its dimensions of 130Å, 1.7 µm, with measurements of 2.1 mm X 100 mm, within a singular package. In regard to the mobile phases, mobile phase A was designated as formic acid at a concentration of 0.1% (v/v) in water, whereas mobile phase B consisted of acetonitrile supplemented with 0.1% (v/v) formic acid. The elution profile unfurled in the following manner: during the interval of 0-0.5 minutes, mobile phase B experienced variation, fluctuating between 3% and 15%; within the time frame of 0.5-2.5 minutes, the concentration of mobile phase B ranged from 15% to 85%; from 2.5-2.6 minutes, a transition took place, escalating from 85% to 100% B; at 2.6-3.5 minutes, mobile phase B remained entirely at 100%; during 3.5-3.6 minutes, a gradual transition ensued from 100% B to 3% B; and finally, from 3.6-6.0 minutes, a consistent level of 3% B was maintained. Noteworthy care was taken in establishing the mass spectrum parameters for the analysis, encompassing an ion spray voltage of 5500 V, a turbine gas temperature of 650°C, a collision gas set at 3 psi, a maintained curtain gas at 35 psi, and an ion source gas pressure amounting to 60 psi. The analysis was conducted under the positive ion mode, and the detection of parent ion product ions was accomplished through the implementation of multiple reaction monitoring (MRM), specifically targeting m/z: 308.1-179.1. The collision energy (CE) was set at 17V, while the declustering potential (DP) was adjusted to 46V.

### RNA-seq transcriptomic analysis

The roots and siliques of WT and *Bnastop2* mutants were collected to facilitate the RNA-seq analysis, with three biological replicates. The extraction of total RNA was conducted employing the TIANGEN RNAprep Pure Plant Kit. An amount of 1.5 μg of RNA per sample was utilized as the input material for the preparation of RNA samples. Subsequently, sequencing libraries were fabricated using the NEBNext® UltraTM RNA Library Prep Kit for Illumina® (NEB, USA). The resulting libraries underwent sequencing on an Illumina Hiseq 4000 platform, which produced 150 bp paired-end reads. The FastQC software was employed for quality control assessment of the RNA-seq reads, and the ensuing results were consolidated using multi-qc (Ewels et al., 2016). We employed the hisat2 algorithm for quantifying the RNA sequencing reads of annotated genes in the Brassica napus genome. The annotated gene data was downloaded from the following website: http://www.genoscope.cns.fr/brassicanapus/. Subsequently, we imported the data and performed normalization using StringTie (Pertea et al., 2015), and DESeq2 was used for differential expression analysis (Love et al., 2014). GO enrichment analysis and KEGG enrichment analysis were conducted using the clusterProfiler package (https://github.com/GuangchuangYu/clusterProfiler).

### Pathogen inoculation and disease resistance assay

The *Sclerotinia sclerotiorum* strain 1980 was cultivated on potato dextrose agar (PDA) medium, which comprised 20% (w/v) potato, 2% (w/v) dextrose, and 1.5% (w/v) agar, for the purpose of inoculation. The leaf inoculation assay was conducted following previously described methods (Zhang et al., 2022), utilizing the latest fully unfolded leaves of 4-week-old plants. After 48 hours of inoculation, photographs of the inoculated leaves were captured, and the diameter of lesions was quantified using a Vernier caliper.

## Supplemental Data

**Supplemental Figure 1.** Comparison of amino acid sequence between STOP1 and STOP2 in Arabidoosis.

**Supplemental Figure 2.** Analysis of the amino acid sequence of *Bnastop2 11-7* mutants.

**Supplemental Figure 3.** Relative RNA level of *BnaSTOP2s* in WT and *Bnastop2* mutants.

**Supplemental Figure 4.** Screening point-to-point validation of BnaA02.1.STOP2 Y2H library.

**Supplemental Figure 5.** Analysis of differential metabolites in roots of WT and *Bnastop2* mutants.

**Supplemental Figure 6.** Intact GSLs identified using the negative ESI-MS/MS mode.

**Supplemental Figure 7.** Seed protein content (%) in WT and *Bnastop2* mutants.

**Supplemental Figure 8.** Determination and analysis of GSH content (ng/ml) in WT and *Bnastop2* mutants.

**Supplemental Figure 9.** Relative RNA level of *BnaMYB28s* in WT and *Bnastop2* mutants.

**Supplemental Figure 10.** Analysis with differentially expressed gene set in roots of *Bnastop2* mutant and WT at 20 DAF.

**Supplemental Figure 11.** Analysis with differentially expressed genes in siliques of *Bnastop2* mutants and WT at 20 DAF.

**Supplemental Figure 12.** Identification of resistance to *Sclerotinia sclerotiorum* on leaves of *Bnastop2* mutants.

**Supplemental Figure 13.** Relative expression of *BnaA02.1.STOP2* in *BnaSTOP2-OE* mutants.

**Supplemental Figure 14.** Analysis with differentially expressed genes of *BnaSTOP2-OE* mutants at 0 hpi and 48 hpi.

**Supplemental Dataset 1.** Detection of mutations at putative CRISPR/Cas9 off-target sites in the T_1_ generation.

**Supplemental Dataset 2.** Relative *BnaSTOP2s* RNA level in WT and *Bnastop2* mutants.

**Supplemental Dataset 3.** N content (mg/L) in the roots and shoots of the *Bnastop2* mutants.

**Supplemental Dataset 4.** Identification of interacting proteins with BnaA02.1.STOP2 through yeast two-hybrid screening.

**Supplemental Dataset 5.** Metabolite profiling of *Bnastop2* mutant roots through LC-MS analysis.

**Supplemental Dataset 6.** The content of GSLs (ng/ml) in WT and *Bnastop2* mutants.

**Supplemental Dataset 7.** The content of GSH (ng/ml) in WT and *Bnastop2* mutants.

**Supplemental Dataset 8.** Relative *BnaMYB28s* and *BnaGTR2s* RNA level in WT and *Bnastop2* mutants.

**Supplemental Dataset 9.** Comparison of protein content (%) in mature seeds between *Bnastop2* mutants and their control in the T_2_ generation.

**Supplemental Dataset 10.** KEGG analysis of up-regulated genes in roots of *Bnastop2* mutants and WT.

**Supplemental Dataset 11.** KEGG analysis of down-regulated genes in roots of *Bnastop2* mutants and WT.

**Supplemental Dataset 12.** KEGG analysis of differentially expressed genes in siliques of *Bnastop2* mutants and WT.

**Supplemental Dataset 13.** DEGs of sulfur metabolism related genes in roots of *Bnastop2* mutants and WT.

**Supplemental Dataset 14.** DEGs of glucosinolate metabolism related genes in roots of *Bnastop2* mutants and WT.

**Supplemental Dataset 15.** DEGs of glucosinolate metabolism related genes in siliques of *Bnastop2* mutants and WT.

**Supplemental Dataset 16.** KEGG analysis of up-regulated genes in *BnaSTOP2-OE* mutants and Xiaoyun (48 hpi).

**Supplemental Dataset 17.** Primers used in this study.

## ACKNOWLEDGEMENTS

This research was supported by the National Key Research and Development Program of China (2022YFD1200400) and the Program for Modern Agricultural Industrial Technology System (CARS-12).

## AUTHOR CONTRIBUTIONS

D.H. and G.Y. designed and supervised the study. L.D., Z.X., T.A. and X.L. performed the bioinformatics analysis. L.D., Z.X., A.L. and J.Z. performed the *STOP2*-related experiments. L.D. and Z.X. prepared the manuscript. D.H. and G.Y. revised the manuscript. All the authors have read and approved the manuscript.

## References

[1]. Aarabi, F., Naake, T., Fernie, A.R., and Hoefgen, R. (2020). Coordinating Sulfur Pools under Sulfate Deprivation. Trends Plant Sci 25: 1227–1239.

[2]. Abuyusuf, M., Robin, A.H.K., Lee, J.H., Jung, H.J., Kim, H.T., Park, J.I., and Nou, I.S. (2018). Glucosinolate Profiling and Expression Analysis of Glucosinolate Biosynthesis Genes Differentiate White Mold Resistant and Susceptible Cabbage Lines. Int J Mol Sci 19.

[3]. Ali, M.A., Azeem, F., Nawaz, M.A., Acet, T., Abbas, A., Imran, Q.M., Shah, K.H., Rehman, H.M., Chung, G., Yang, S.H., and Bohlmann, H. (2018). Transcription factors WRKY11 and WRKY17 are involved in abiotic stress responses in Arabidopsis. J Plant Physiol 226: 12–21.

[4]. Brosnan, J.T., and Brosnan, M.E. (2006). The sulfur-containing amino acids: An overview. J Nutr 136: 1636s–1640s.

[5]. Burow, M., Zhang, Z.Y., Ober, J.A., Lambrix, V.M., Wittstock, U., Gershenzon, J., and Kliebenstein, D.J. (2008). ESP and ESM1 mediate indol-3-acetonitrile production from indol-3-ylmethyl glucosinolate in Arabidopsis. Phytochemistry 69: 663–671.

[6]. Buxdorf, K., Yaffe, H., Barda, O., and Levy, M. (2013). The Effects of Glucosinolates and Their Breakdown Products on Necrotrophic Fungi. Plos One 8.

[7]. Capaldi, F.R., Gratao, P.L., Reis, A.R., Lima, L.W., and Azevedo, R.A. (2015). Sulfur Metabolism and Stress Defense Responses in Plants. Trop Plant Biol 8: 60–73.

[8]. Chan, K.X., Phua, S.Y., and Van Breusegem, F. (2019). Secondary sulfur metabolism in cellular signalling and oxidative stress responses. J Exp Bot 70: 4237–4250.

[9]. Chao, H.B., Wang, H., Wang, X.D., Guo, L.X., Gu, J.W., Zhao, W.G., Li, B.J., Chen, D.Y., Raboanatahiry, N., and Li, M.T. (2017). Genetic dissection of seed oil and protein content and identification of networks associated with oil content in Brassica napus. Sci Rep-Uk 7.

[10]. Chen, H.M., Zou, Y., Shang, Y.L., Lin, H.Q., Wang, Y.J., Cai, R., Tang, X.Y., and Zhou, J.M. (2008). Firefly luciferase complementation imaging assay for protein-protein interactions in plants. Plant Physiol 146: 368–376.

[11]. Chen, J.Y., Ullah, C., Reichelt, M., Beran, F., Yang, Z.L., Gershenzon, J., Hammerbacher, A., and Vassao, D.G. (2020). The phytopathogenic fungus Sclerotinia sclerotiorum detoxifies plant glucosinolate hydrolysis products via an isothiocyanate hydrolase. Nat Commun 11.

[12]. Chittem, K., Yajima, W.R., Goswami, R.S., and Mendoza, L.E.D. (2020). Transcriptome analysis of the plant pathogen Sclerotinia sclerotiorum interaction with resistant and susceptible canola (Brassica napus) lines. Plos One 15.

[13]. Ewels, P., Magnusson, M., Lundin, S., and Kaller, M. (2016). MultiQC: summarize analysis results for multiple tools and samples in a single report. Bioinformatics 32: 3047–3048.

[14]. Francioso, A., Conrado, A.B., Mosca, L., and Fontana, M. (2020). Chemistry and Biochemistry of Sulfur Natural Compounds: Key Intermediates of Metabolism and Redox Biology. Oxid Med Cell Longev 2020.

[15]. Fujiwara, T., Nambara, E., Yamagishi, K., Goto, D.B., and Naito, S. (2002). Storage proteins. Arabidopsis Book 1: e0020.

[16]. Gan, L., Sun, X., Jin, L., Wang, G., Xu, J., Wei, Z., and Fu, T. (2003). Establishment of math models of NIRS analysis for oil and protein contents in seed ofo Brassica napus. Zhongguo Nong Ye Ke Xue 36: 1609–1613.

[17]. Grant, C.A., Mahli, S.S., and Karamanos, R.E. (2012). Sulfur management for rapeseed. Field Crop Res 128: 119–128.

[18]. Hatzfeld, Y., Lee, S., Lee, M., Leustek, T., and Saito, K. (2000). Functional characterization of a gene encoding a fourth ATP sulfurylase isoform from Arabidopsis thaliana. Gene 248: 51–58.

[19]. Hawkesford, M.J. (2012). Sulfate Uptake and Assimilation – Whole Plant Regulation Dordrecht: Springer Netherlands. 11–24.

[20]. Hawkesford, M.J., and De Kok, L.J. (2006). Managing sulphur metabolism in plants. Plant Cell Environ 29: 382–395.

[21]. Heeg, C., Kruse, C., Jost, R., Gutensohn, M., Ruppert, T., Wirtz, M., and Hell, R. (2008). Analysis of the Arabidopsis O-acetylserine(thiol)lyase gene family demonstrates compartment-specific differences in the regulation of cysteine synthesis. Plant Cell 20: 168–185.

[22]. Hunziker, P., Ghareeb, H., Wagenknecht, L., Crocoll, C., Halkier, B.A., Lipka, V., and Schulz, A. (2020). De novo indol-3-ylmethyl glucosinolate biosynthesis, and not long-distance transport, contributes to defence of Arabidopsis against powdery mildew. Plant Cell Environ 43: 1571–1583.

[23]. Ito, T., Kitaiwa, T., Nishizono, K., Umahashi, M., Miyaji, S., Agake, S.I., Kuwahara, K., Yokoyama, T., Fushinobu, S., Maruyama-Nakashita, A., Sugiyama, R., Sato, M., Inaba, J., Hirai, M.Y., and Ohkama-Ohtsu, N. (2022). Glutathione degradation activity of gamma-glutamyl peptidase 1 manifests its dual roles in primary and secondary sulfur metabolism in Arabidopsis. Plant J 111: 1626–1642.

[24]. Jez, J.M. (2019). Structural biology of plant sulfur metabolism: from sulfate to glutathione. J Exp Bot 70: 4089–4103.

[25]. Jiang, C.H., Huang, Z.Y., Xie, P., Gu, C., Li, K., Wang, D.C., Yu, Y.Y., Fan, Z.H., Wang, C.J., Wang, Y.P., Guo, Y.H., and Guo, J.H. (2016). Transcription factors WRKY70 and WRKY11 served as regulators in rhizobacterium Bacillus cereus AR156-induced systemic resistance to Pseudomonas syringae pv. tomato DC3000 in Arabidopsis. J Exp Bot 67: 157–174.

[26]. Joshi, N.C., Meyer, A.J., Bangash, S.A.K., Zheng, Z.L., and Leustek, T. (2019). Arabidopsis γ-glutamylcyclotransferase affects glutathione content and root system architecture during sulfur starvation. New Phytol 221: 1387–1397.

[27]. Kamal, M.M., Savocchia, S., Lindbeck, K.D., and Ash, G.J. (2016). Biology and biocontrol of Sclerotinia sclerotiorum (Lib.) de Bary in oilseed Brassicas. Australas Plant Path 45: 1–14.

[28]. Kobayashi, Y., Ohyama, Y., Kobayashi, Y., Ito, H., Iuchi, S., Fujita, M., Zhao, C.R., Tanveer, T., Ganesan, M., Kobayashi, M., and Koyama, H. (2014). STOP2 Activates Transcription of Several Genes for Al- and Low pH-Tolerance that Are Regulated by STOP1 in Arabidopsis. Mol Plant 7: 311–322.

[29]. Kopriva, S., Malagoli, M., and Takahashi, H. (2019). Sulfur nutrition: impacts on plant development, metabolism, and stress responses. J Exp Bot 70: 4069–4073.

[30]. Li-Beisson, Y., Beisson, F., and Riekhof, W. (2015). Metabolism of acyl-lipids in Chlamydomonas reinhardtii. Plant J 82: 504–522.

[31]. Livak, K.J., and Schmittgen, T.D. (2001). Analysis of relative gene expression data using real-time quantitative PCR and the 2(T)(-Delta Delta C) method. Methods 25: 402–408.

[32]. Love, M.I., Huber, W., and Anders, S. (2014). Moderated estimation of fold change and dispersion for RNA-seq data with DESeq2. Genome Biol 15: 550.

[33]. Maruyama-Nakashita, A., Nakamura, Y., Tohge, T., Saito, K., and Takahashi, H. (2006). SLIM1 is a central transcriptional regulator of plant sulfur response and metabolism. Plant Cell 18: 3235–3251.

[34]. Miller, A.J., and Chapman, N. (2011). Transporters Involved in Nitrogen Uptake and Movement. In: The Molecular and Physiological Basis of Nutrient Use Efficiency in Crops. 193–210.

[35]. Mugford, S.G., Matthewman, C., Lee, B.-R., Yatusevich, R., Yoshimoto, N., Wirtz, M., Hill, L., Hell, R., Takahashi, H., Saito, K., Gigolashvili, T., and Kopriva, S. (2012). Partitioning of Sulfur Between Primary and Secondary Metabolism Dordrecht: Springer Netherlands. 91–96.

[36]. Mugford, S.G., Yoshimoto, N., Reichelt, M., Wirtz, M., Hill, L., Mugford, S.T., Nakazato, Y., Noji, M., Takahashi, H., Kramell, R., Gigolashvili, T., Flugge, U.I., Wasternack, C., Gershenzon, J., Hell, R., Saito, K., and Kopriva, S. (2009). Disruption of Adenosine-5 ’-Phosphosulfate Kinase in Arabidopsis Reduces Levels of Sulfated Secondary Metabolites. Plant Cell 21: 910–927.

[37]. Narayan, O.P., Kumar, P., Yadav, B., Dua, M., and Johri, A.K. (2022). Sulfur nutrition and its role in plant growth and development. Plant Signal Behav.

[38]. Noman, A., Aqeel, M., Khalid, N., Islam, W., Sanaullah, T., Anwar, M., Khan, S., Ye, W.F., and Lou, Y.G. (2019). Zinc finger protein transcription factors: Integrated line of action for plant antimicrobial activity. Microb Pathogenesis 132: 141–149.

[39]. Nour-Eldin, H.H., Andersen, T.G., Burow, M., Madsen, S.R., Jorgensen, M.E., Olsen, C.E., Dreyer, I., Hedrich, R., Geiger, D., and Halkier, B.A. (2012). NRT/PTR transporters are essential for translocation of glucosinolate defence compounds to seeds. Nature 488: 531–534.

[40]. Paulose, B., Chhikara, S., Coomey, J., Jung, H.I., Vatamaniuk, O., and Dhankher, O.P. (2013). A gamma-Glutamyl Cyclotransferase Protects Arabidopsis Plants from Heavy Metal Toxicity by Recycling Glutamate to Maintain Glutathione Homeostasis. Plant Cell 25: 4580–4595.

[41]. Pertea, M., Pertea, G.M., Antonescu, C.M., Chang, T.C., Mendell, J.T., and Salzberg, S.L. (2015). StringTie enables improved reconstruction of a transcriptome from RNA-seq reads. Nat Biotechnol 33: 290-+.

[42]. Petersen, B.L., Chen, S.X., Hansen, C.H., Olsen, C.E., and Halkier, B.A. (2002). Composition and content of glucosinolates in developing Arabidopsis thaliana. Planta 214: 562–571.

[43]. Purnamasari, M., Cawthray, G.R., Barbetti, M.J., Erskine, W., and Croser, J.S. (2015). Camalexin Production in Camelina sativa is Independent of Cotyledon Resistance to Sclerotinia sclerotiorum. Plant Dis 99: 1544–1549.

[44]. Rahikainen, M., Trotta, A., Alegre, S., Pascual, J., Vuorinen, K., Overmyer, K., Moffatt, B., Ravanel, S., Glawischnig, E., and Kangasjarvi, S. (2017). PP2A-B ’gamma modulates foliar trans-methylation capacity and the formation of 4-methoxy-indol-3-yl-methyl glucosinolate in Arabidopsis leaves. Plant J 89: 112–127.

[45]. Samizadeh, H., Yazdi-samadi, B., Bihamta, M.R., Taleii, A.R., and Stringam, G.R. (2007). Study of Pod Length Trait in Doubled Haploid Brassica napus Population by Molecular Markers. J Agr Sci Tech-Iran 9: 129–136.

[46]. Sanden, N.C.H., Kanstrup, C., Crocoll, C., Schulz, A., Nour-Eldin, H.H., Halkier, B.A., and Xu, D.Y. (2024). An UMAMIT-GTR transporter cascade controls glucosinolate seed loading in. Nat Plants.

[47]. Setya, A., Murillo, M., and Leustek, T. (1996). Sulfate reduction in higher plants: Molecular evidence for a novel 5’-adenylylsulfate reductase. P Natl Acad Sci USA 93: 13383–13388.

[48]. Sotelo, T., Lema, M., Soengas, P., Cartea, M.E., and Velasco, P. (2015). In Vitro Activity of Glucosinolates and Their Degradation Products against Brassica-Pathogenic Bacteria and Fungi. Appl Environ Microb 81: 432–440.

[49]. Stotz, H.U., Sawada, Y., Shimada, Y., Hirai, M.Y., Sasaki, E., Krischke, M., Brown, P.D., Saito, K., and Kamiya, Y. (2011). Role of camalexin, indole glucosinolates, and side chain modification of glucosinolate-derived isothiocyanates in defense of Arabidopsis against Sclerotinia sclerotiorum. Plant J 67: 81–93.

[50]. Subramanian, A., Tamayo, P., Mootha, V.K., Mukherjee, S., Ebert, B.L., Gillette, M.A., Paulovich, A., Pomeroy, S.L., Golub, T.R., Lander, E.S., and Mesirov, J.P. (2005). Gene set enrichment analysis: A knowledge-based approach for interpreting genome-wide expression profiles. P Natl Acad Sci USA 102: 15545–15550.

[51]. Takahashi, H., Kopriva, S., Giordano, M., Saito, K., and Hell, R. (2011). Sulfur Assimilation in Photosynthetic Organisms: Molecular Functions and Regulations of Transporters and Assimilatory Enzymes. Annual Review of Plant Biology, Vol 62 62: 157–184.

[52]. Tan, Z., Xie, Z., Dai, L., Zhang, Y., Hu, Z., Tang, S., Wan, L., Yao, X., Guo, L., and Hong, D. (2021). Genome- and transcriptome-wide association studies reveal the genetic basis and the breeding history of seed glucosinolate content in Brassica napus. Plant Biotechnol J.

[53]. Teng, Z.Y., Yu, Y.J., Zhu, Z.J., Hong, S.B., Yang, B.X., and Zang, Y.X. (2021). Melatonin elevated Sclerotinia sclerotiorum resistance via modulation of ATP and glucosinolate biosynthesis in Brassica rapa ssp. pekinensis. J Proteomics 243.

[54]. Tokizawa, M., Enomoto, T., Chandnani, R., Mora-Macías, J., Burbridge, C., Armenta-Medina, A., Kobayashi, Y., Yamamoto, Y.Y., Koyama, H., and Kochian, L.V. (2023). The transcription factors, STOP1 and TCP20, are required for root system architecture alterations in response to nitrate deficiency. Proceedings of the National Academy of Sciences 120: e2300446120.

[55]. Vauclare, P., Kopriva, S., Fell, D., Suter, M., Sticher, L., von Ballmoos, P., Krahenbuhl, U., den Camp, R.O., and Brunold, C. (2002). Flux control of sulphate assimilation in Arabidopsis thaliana: adenosine 5 ’-phosphosulphate reductase is more susceptible than ATP sulphurylase to negative control by thiols. Plant J 31: 729–740.

[56]. Venditti, A., and Bianco, A. (2020). Sulfur-containing Secondary Metabolites as Neuroprotective Agents. Curr Med Chem 27: 4421–4436.

[57]. Verma, S., Singh, A., Pradhan, S., Kumar, V., and Kumar, V. (2022). Effect of Sulphur Nutrition on the Production Potential of Brassica spp.: A Review. 880–887.

[58]. Wang, B., Wu, Z.K., Li, Z.H., Zhang, Q.H., Hu, J.L., Xiao, Y.J., Cai, D.F., Wu, J.S., King, G.J., Li, H.T., and Liu, K.D. (2018). Dissection of the genetic architecture of three seed-quality traits and consequences for breeding in Brassica napus. Plant Biotechnol J 16: 1336–1348.

[59]. Wirtz, M., Droux, M., and Hell, R. (2004). O-acetylserine (thiol) lyase: an enigmatic enzyme of plant cysteine biosynthesis revisited in Arabidopsis thaliana. J Exp Bot 55: 1785–1798.

[60]. Wirtz, M., Heeg, C., Samami, A.A., Ruppert, T., and Hell, R. (2010). Enzymes of cysteine synthesis show extensive and conserved modifications patterns that include N-alpha-terminal acetylation. Amino Acids 39: 1077–1086.

[61]. Xing, H.L., Dong, L., Wang, Z.P., Zhang, H.Y., Han, C.Y., Liu, B., Wang, X.C., and Chen, Q.J. (2014). A CRISPR/Cas9 toolkit for multiplex genome editing in plants. Bmc Plant Biol 14.

[62]. Xu, D.Y., Sanden, N.C.H., Hansen, L.L., Belew, Z.M., Madsen, S.R., Meyer, L., Jorgensen, M.E., Hunziker, P., Veres, D., Crocoll, C., Schulz, A., Nour-Eldin, H.H., and Halkier, B.A. (2023). Export of defensive glucosinolates is key for their accumulation in seeds. Nature 617: 132-+.

[63]. Yang, L.Y., Zhang, Y., Guan, R.X., Li, S., Xu, X.W., Zhang, S.Q., and Xu, J. (2020). Co-regulation of indole glucosinolates and camalexin biosynthesis by CPK5/CPK6 and MPK3/MPK6 signaling pathways. J Integr Plant Biol 62: 1780–1796.

[64]. Ye, J.Y., Tian, W.H., Zhou, M., Zhu, Q.Y., Du, W.X., Zhu, Y.X., Liu, X.X., Lin, X.Y., Zheng, S.J., and Jin, C.W. (2021). STOP1 activates NRT1.1-mediated nitrate uptake to create a favorable rhizospheric pH for plant adaptation to acidity. Plant Cell 33: 3658–3674.

[65]. Yu, B.Y., Gruber, M., Khachatourians, G.G., Hegedus, D.D., and Hannoufa, A. (2010). Gene expression profiling of developing Brassica napus seed in relation to changes in major storage compounds. Plant Sci 178: 381–389.

[66]. Zhang, K., Liu, F., Wang, Z.X., Zhuo, C.J., Hu, K.N., Li, X.X., Wen, J., Yi, B., Shen, J.X., Ma, C.Z., Fu, T.D., and Tu, J.X. (2022). Transcription factor WRKY28 curbs WRKY33-mediated resistance to Sclerotinia sclerotiorum in Brassica napus. Plant Physiol 190: 2757–2774.

[67]. Zhang, L., Kawaguchi, R., Morikawa-Ichinose, T., Allahham, A., Kim, S.J., and Maruyama-Nakashita, A. (2020). Sulfur Deficiency-Induced Glucosinolate Catabolism Attributed to Two beta-Glucosiaases, BGLU28 and BGLU30, is Required for Plant Growth Maintenance under Sulfur Deficiency. Plant Cell Physiol 61: 803–813.

[68]. Zhao, F.-j., Tausz, M., and De Kok, L.J. (2008). Role of Sulfur for Plant Production in Agricultural and Natural Ecosystems. In: Sulfur Metabolism in Phototrophic Organisms--Hell, R., Dahl, C., Knaff, D., and Leustek, T., eds. Dordrecht: Springer Netherlands. 417–435.

[69]. Zhao, Q., Wu, Y., Gao, L., Ma, J., Li, C.Y., and Xiang, C.B. (2014). Sulfur nutrient availability regulates root elongation by affecting root indole-3-acetic acid levels and the stem cell niche. J Integr Plant Biol 56: 1151–1163.

[70]. Zhou, X.M., Zhang, H.Y., Xie, Z.Q., Liu, Y., Wang, P.F., Dai, L.H., Zhang, X.H., Wang, Z.Y., Wang, Z.A.R., Wan, L.L., Yang, G.S., and Hong, D.F. (2023). Natural variation and artificial selection at the BnaC2.MYB28 locus modulate Brassica napus seed glucosinolate. Plant Physiol 191: 352–368.

